# Mapping the Human Visual Thalamus and its Cytoarchitectonic Subdivisions Using Quantitative MRI

**DOI:** 10.1101/2020.08.14.250779

**Authors:** Christa Müller-Axt, Cornelius Eichner, Henriette Rusch, Louise Kauffmann, Pierre-Louis Bazin, Alfred Anwander, Markus Morawski, Katharina von Kriegstein

## Abstract

The human lateral geniculate nucleus (LGN) of the visual thalamus is a key subcortical processing site for visual information analysis. A non-invasive assessment of the LGN and its functionally and microstructurally distinct magnocellular (M) and parvocellular (P) subdivisions *in-vivo* in humans is challenging, because of its small size and location deep inside the brain. Here we tested whether recent advances in high-field structural quantitative MRI (qMRI) can enable MR-based mapping of human LGN subdivisions. We employed ultra-high resolution 7 Tesla qMRI of a *post-mortem* human LGN specimen and high-resolution 7 Tesla *in-vivo* qMRI in a large participant sample. We found that a quantitative assessment of the LGN and a differentiation of its subdivisions was possible based on microstructure-informed MR-contrast alone. In both the *post-mortem* and *in-vivo* qMRI data, we identified two components of shorter and longer longitudinal relaxation time (T_1_) within the LGN that coincided with the known anatomical locations of a dorsal P and a ventral M subdivision, respectively. Through a subsequent ground truth histological examination of the same *post-mortem* LGN specimen, we showed that the observed T_1_ contrast pertains to cyto- and myeloarchitectonic differences between LGN subdivisions. These differences were based on cell and myelin density, but not on iron content. Our qMRI-based mapping strategy overcomes shortcomings of previous fMRI-based mapping approaches. It paves the way for an in-depth understanding of the function and microstructure of the LGN in humans. It also enables investigations into the selective contributions of LGN subdivisions to human behavior in health and disease.

**Significance Statement:** The lateral geniculate nucleus (LGN) is a key processing site for the analysis of visual information. Due to its small size and deep location within the brain, non-invasive mapping of the LGN and its microstructurally distinct subdivisions in humans is challenging. Using quantitative MRI methods that are sensitive to underlying microstructural tissue features, we show that a differentiation of the LGN and its microstructurally distinct subdivisions is feasible in humans non-invasively. These findings are important because they open up novel opportunities to assess the hitherto poorly understood complex role of the LGN in human perception and cognition, as well as the contribution of selective LGN subdivision impairments to various clinical conditions including developmental dyslexia, glaucoma and multiple sclerosis.

## Introduction

The human sensory thalami are central processing sites for the analysis of sensory information. A growing body of empirical evidence suggests that to-date we are only starting to understand the function of these nuclei and their subdivisions for human behavior and cognition in health (1) and disease (2–5). One reason for this lack of understanding is due to severe technical difficulties in assessing these nuclei *in-vivo* in humans using non-invasive conventional magnetic resonance imaging (MRI) (6). Here, we show that the visual lateral geniculate nucleus (LGN) and its cytoarchitectonic subdivisions can be identified in humans both *in-vivo* and *post-mortem* using advanced microstructure-informed quantitative MRI strategies. This opens up the possibility to use MRI as a tool for an in-depth understanding of the LGN and could be the basis for a future diagnostic tool for LGN integrity in disease.

Most of our current knowledge about the LGN stems from electrophysiological and lesion studies in nonhuman primates (7). While these research findings have greatly contributed to our current understanding of LGN function, solely depending on insights from primate research entails two inherent problems: (i) due to species differences direct inferences from animal studies to humans cannot readily be made (8), and (ii) research suggests that the integrity of subcortical sensory brain structures is relevant to explain human-specific behavior such as human communication (9–11). Overcoming technical issues related to *in-vivo* MR acquisitions of the LGN and its functionally distinct subdivisions will, therefore, be necessary to obtain a complete account of LGN function and dysfunction in humans.

The human LGN consists of six distinct neuronal layers, two ventral magnocellular (M) layers, four dorsal parvocellular (P) layers, and intercalated koniocellular (K) layers (12). The investigation of the LGN’s main neuronal layers (often coined as M and P subdivisions) in humans *in-vivo* faces considerable challenges. First, the LGN’s small size, ranging from only 91-157 mm^3^ in humans (12), and deep location within the brain makes it difficult to map the LGN using non-invasive MR imaging techniques (6). Second, conventional image resolutions in *in-vivo* MR examinations (structural MRI ~1mm, functional MRI ~2.5mm isotropic resolution) are likely insufficient to fully disentangle distinct signal contributions of M and P subdivisions due to partial volume effects (13). Third, any microstructural tissue differences between LGN subdivisions can be assumed to result in only subtle differences in MR contrast, which further complicates their differentiation (i.e., all LGN subdivisions are composed of subcortical gray matter, whereas MRI contrast is strongest between different tissue types, like white- and gray matter).

Resolving LGN subdivisions in humans *in-vivo* would, however, be of keen interest to gain a better understanding of their contribution to a variety of cognitive processes such as visual perception, selective attention and visual awareness (14–16); their functional interactions as well as structural connectivity patterns with the cerebral cortex (17); and their role in human disorders such as glaucoma (18, 19), multiple sclerosis (20, 21), developmental dyslexia (2, 3, 22), autism spectrum- (23) and mood disorders (4). Indeed, most clinical research on the integrity of the LGN and its subdivisions in humans is based on *post-mortem* studies, which generally lack the opportunity to relate structure to function and often suffer from small sample sizes. They, however, also give first indications that LGN subdivisions can be selectively impaired in clinical disorders, emphasizing the need for a sub-parcellation of the LGN (3, 4, 20).

Recent advances in high-field structural MRI have enabled measurements at an unprecedented level of spatial resolution. Especially the introduction of quantitative MRI (qMRI) methods provides access to imaging data with improved image contrast, which also allows researchers to assess biophysical tissue parameters that reflect the underlying microstructure (24). Quantitative MR parameters, such as the longitudinal relaxation time T_1_, provide insights into the degree of tissue myelination per voxel (25–27) and allow tissue segmentation based on cyto- and myeloarchitecture (28–30). To our knowledge, no study has yet attempted to resolve LGN subdivisions *in-vivo* based on microstructure-informed contrasts in structural MRI. However, such an approach seems promising as M and P layers differ markedly in cyto- and myeloarchitecture. In primates, including humans, M neurons are characterized by large cell bodies and are more sparsely distributed relative to the smaller, densely packed P neurons (7, 12, 31, 32). In addition, M neurons have thicker and more myelinated nerve fibers enabling faster signal transmissions compared to the thinner, less myelinated axons of P neurons (32–37). Mapping strategies based on structural MRI could offer a way to mitigate limitations of previous attempts to resolve LGN subdivisions in humans *in-vivo.* Using extensive functional MRI (fMRI) protocols, a few recent studies identified M and P-like activation clusters within the human LGN by exploiting the distinct functional response sensitivities of M and P neurons (19, 38, 39). While these studies have provided first promising insights into the feasibility of MR-based mapping of LGN subdivisions *in-vivo*, fMRI acquisitions typically suffer from several technical shortcomings such as limited image resolution, blurring, geometric distortions, and lengthy acquisition times (40).

Here, we utilized recent technological advances in high-field structural qMRI (24) and addressed whether microstructure-informed MR contrasts alone can enable MR-based mapping of human LGN subdivisions. The study involved three steps: (i) ultra-high resolution 7 Tesla qMRI of a *post-mortem* human LGN specimen (220μm isotropic) (ii) histology of the same LGN specimen, and (iii) *in-vivo* high-resolution 7 Tesla qMRI assessment of N=27 bilateral LGNs (500μm isotropic). We expected that microstructural tissue differences between LGN subdivisions are reflected in T_1_ relaxation and that this subdivisional T_1_ contrast is driven by local differences in myelin density between M and P layers. As myelin density and T_1_ relaxation are inversely related (25, 27), a key question is which cyto- and myeloarchitectonic features of the LGN primarily constitute myelin density and, thus, T_1_ contrast between LGN subdivisions. First, higher axonal myelination of M than P neurons (34–36) might result in an increased myelin density, and consequently, shorter T_1_ relaxation of the M relative to the P subdivision. Alternatively, the overall sparser distribution of M neurons (7, 12) might decrease myelin density and translate into increased T_1_ relaxation of the M relative to the P subdivision. To validate and explain potential quantitative T_1_ contrasts between LGN subdivisions, we compared our qMRI results to T_1_-relevant measures obtained from histology. With the *in-vivo* data, we aimed to extend our *post-mortem* qMRI analyses to a broader scientific and clinical purpose. The *in-vivo* data were also used to create a detailed population atlas of the LGN and its M and P subdivisions, which we have made publicly available.

## Material and Methods

### *Post-Mortem* MRI and Histology

#### Collection and Preparation of Post-Mortem Human Brain Tissue

A human *post-mortem* brain sample was provided by the former Brain Banking Centre Leipzig of the German Brain-Net, operated by the Paul Flechsig Institute of Brain Research, Medical Faculty, University of Leipzig. The brain sample consisted of a left hemisphere of a female patient (89yrs, cause of death myocardial infarction, tissue fixation 24h *post-mortem*). Neuropathological assessment revealed no signs of any neurological diseases. The entire procedure of case recruitment, acquisition of the patient’s personal data, protocols and informed consent forms, performing the autopsy, and handling the autopsy material has been approved by the responsible authorities (GZ 01GI9999-01GI0299; Approval # 82-02). Following standard Brain Bank procedures, the block was immersion-fixed in 3% paraformaldehyde and 1% glutaraldehyde in phosphate-buffered saline (PBS; pH 7.4) for at least six weeks. The tissue block was cut to approximately 30×15×30mm (left-right, posterior-anterior, superior-inferior dimension, respectively) in size and included the LGN and part of the hippocampus. To prepare the sample for high-resolution qMRI, the LGN specimen was placed in an acrylic sphere of 60 mm diameter filled with perfluoropolyether (PFPE; Fomblin©, Solvay Solexis Inc., Bollate, Italy). PFPE is a synthetic oil that does not generate any MR signal. Its application, therefore, has the specific advantage that the measured *post-mortem* qMR parameters are not affected by partial volume effects of other signal-generating substances (41) adjacent to the analyzed tissue.

#### Quantitative MRI Data Acquisition and Reconstruction

For the qMRI acquisition of the LGN specimen, we employed a multi-contrast steady-state approach. The basic principle of this method is to acquire 3D FLASH MR data with different contrasts, which are subsequently fit to an MR signal model (42). As a result, quantitative maps are obtained for T_1_ and proton density (PD). These maps describe physical quantities and are thus described with the prefix ‘q’ to avoid potential confusion with commonly employed non-quantitative contrasts, such as T_1_-weighted or PD-weighted contrasts.

Ultra-high resolution *post-mortem* MRI data were acquired on a 7T Magnetom MRI system (Siemens Healthineers, Erlangen, Germany) using a custom-built Helmholtz coil (43). The use of a Helmholtz coil is especially beneficial for multi-contrast steady-state acquisitions due to its homogenous excitation and receive field. FLASH data were acquired using the following imaging parameters: 220μm isotropic resolution, TE=4-40.7ms (12 echo times), TR=95ms, FoV=50×50×24.64mm^3^, BW=343Hz/Px, slab-selective RF excitation, no GRAPPA, and PF=8/8. Three MR contrasts were obtained by variation of the excitation flip angle (α): a PD-weighted (PDw) contrast at α_PD_=17°, a high-signal Ernst angle contrast at α_Ernst_=32°, and a T_1_-weighted (T_1_w) contrast at α_T1_=82°. Throughout the MR acquisition, the temperature of the LGN specimen was monitored to remain within tissue-preserving ranges.

Quantitative MR parameters were calculated from the MR imaging data in a two-stage procedure using Python (44). First, the echo times of all contrasts were evaluated jointly to determine voxel-wise values for T_2_*. All FLASH contrasts were then extrapolated to TE=0ms to remove T_2_* contrast contamination (45). In a second step, the extrapolated data of all flip angles were fit voxel-wise to the steady-state Ernst-equation (42) to calculate quantitative maps of qT_1_ and qPD. To remove potential impacts from smooth, low frequency field inhomogeneity, the two quantitative maps were bias-field corrected using N4 inhomogeneity correction in ANTs (46).

#### LGN Segmentation on *Post-Mortem* Quantitative T_1_ Map

The LGN was defined on the bias-field corrected ultra-high resolution qT_1_ map of the *post-mortem* qMRI acquisition through manual segmentation by two independent raters in FSLView (https://fsl.fmrib.ox.ac.uk/fsl/fslwiki/FslView). We used the qT_1_ map for segmentation, as this map provides excellent gray-white matter contrast and therefore allows for better visualization and improved segmentation of deep gray matter structures such as the LGN (2, 47, 48). Windowing parameters for the manual segmentation were chosen to maximize LGN contrast and were identical for both raters. Inter-rater reliability for the manual LGN segmentation was assessed by means of a Dice coefficient (49). The obtained coefficient yields a measure of agreement between raters and ranges from 0 (no agreement) to 1 (perfect agreement). Segmentations of both raters were conjoined to only include those voxels that were segmented by both raters into the LGN mask. The LGN mask was used in subsequent analyses of *post-mortem* LGN subdivisions.

#### LGN Subdivisions in *Post-Mortem* Tissue

For the identification of *post-mortem* LGN subdivisions, we extracted the distribution of qT_1_ values within the LGN mask. An automated data-driven segmentation of *post-mortem* LGN subdivisions was implemented by fitting a mixture model with two components, each representing one of the two main LGN subdivisions, to the distribution of *post-mortem* qT_1_ values in the LGN (SI Methods). The distribution of qT_1_ values was assumed to be Gaussian for each subdivision. The extracted LGN qT_1_ distribution was first cleaned from outliers by removing the 0.1% smallest and 0.1% largest qT_1_ values. A Gaussian mixture model was then fit to the cleaned distribution using the SciPy function ‘curve_fit’ (44). Next, each of the two individual Gaussian fits (i.e., M and P components) were normalized by the envelope of the Gaussian mixture model (i.e., whole distribution of P+M components) to transfer LGN qT_1_ intensities into distribution-based probabilities. The distribution probabilities were then interpolated using the SciPy function ‘interp1d’ to compute a continuous transfer function between qT_1_ intensities and distribution probabilities. In a final step, the transfer function was applied to the ultra-high resolution *post-mortem* LGN qT_1_ map to compute voxel-wise M and P distribution probability maps.

#### Histology

To validate and explain potential qT_1_ contrasts between LGN subdivisions, the *post-mortem* LGN sample was subjected to microstructural histology. We included three types of markers that covered the main microstructural properties contributing to qT1 contrast (27): this included immunohistochemical staining with anti-human neuronal protein C/D (HuC/D) and myelin basic protein (MBP) for marking cell bodies and myelin, respectively. In addition, Perls’ Prussian blue (PB) as a marker for ferric iron was included as another potential source of qT_1_ contrast (27). Detailed procedures for immunohistochemical staining with HuC/D and MBP as well as histochemical staining with PB are described in the supplementary materials (SI Methods). Following (immuno-)histochemical staining, measures of cell density based on anti-HuC/D, myelin density based on MBP optical density, and iron content based on PB optical density were extracted for each of the two fused dorsal P layers (i.e., layers P3/5 and P4/6) and each of the two ventral M layers of the LGN (SI Methods).

### *In-vivo* MRI

#### Participants

To address whether qMRI can also guide the differentiation of LGN subdivisions *in-vivo*, we analyzed high-resolution structural qMRI data from N=27 (15 females, 12 males) healthy participants with a mean age of 26.5±3.8 years. The data were obtained from an open access repository of high-resolution and quantitative MR brain imaging data (50) (available for download at http://openscience.cbs.mpg.de/bazin/7T_Quantitative/). Of the N=28 publicly available datasets in the repository, the dataset of one participant was omitted due to a diagnosis of developmental dyslexia. All participants, except for one, were right-handed as assessed with the Edinburgh Inventory (51), and none had a prior history of neurological or psychiatric disorders. Written informed consent was obtained from all participants. The study was approved by the ethics committee of the Medical Faculty, University of Leipzig, Germany (Approval # 177-2007).

#### High-Resolution 7 Tesla Quantitative MRI Data Acquisition

High-resolution structural qMRI data were acquired on a 7 Tesla Magnetom MRI system (Siemens Healthineers, Erlangen, Germany) using a 24-channel head array coil (NOVA Medical, Wilmington MA, USA). Each hemisphere was imaged separately using a 3D-MP2RAGE sequence (48) with the following imaging parameters: 500μm isotropic resolution, TE=2.45ms, TR=5000ms, TI_1_/TI_2_=900/2750ms, α_1_/α_2_=3/5°, FoV=224×224×104mm^3^, and Partial Fourier (PF) of 6/8 in slice direction. The acquisition took approximately 28 minutes per hemisphere. The 3D-MP2RAGE sequence included two readouts at different inversion times, from which a qT_1_ map was estimated. A single whole-brain qT_1_ map of the two hemispheres was created by co-registering both images and interpolating the result at 400μm isotropic resolution using a rigid (6-parameter) transformation. Finally, the qT_1_ maps of all participants were skull-stripped (28). All processing was performed using CBS Tools (https://www.nitrc.org/projects/cbs-tools/).

#### LGN Segmentations on *In-vivo* Quantitative T_1_ maps

Bilateral LGNs were defined through manual segmentation by two independent raters on the N=27 high-resolution *in-vivo* q_T1_ maps following a standardized procedure in FSLView (SI Methods). Dice coefficients were computed to assess inter-rater reliability. The LGN segmentations of both raters were merged to create an LGN mask for each participant and each hemisphere. For the LGN masks, only those voxels that were segmented by both raters were considered.

#### *In-Vivo* LGN Subdivisions

For the identification of *in-vivo* LGN subdivisions, we first normalized the LGN masks of all participants into a common reference space. This was done by computing a study-specific qT_1_ group template from the N=27 *in-vivo* high-resolution qT_1_ maps using symmetric normalization (SyN) in ANTs (52) (SI Methods and Fig. S1A-C). SyN is a state-of-the-art registration algorithm that globally minimizes registration parametrization and is unbiased towards any individual input image in template generation (53). The obtained SyN registration parameters were applied to the individual qT_1_ maps, followed by averaging of all registered maps, to create the study-specific qT_1_ group template (SI Methods and Fig. S1C). The respective registration parameters were also applied to the individual LGN masks, which were combined for each hemisphere to create a bilateral LGN population atlas in a common reference (i.e., template) space (SI Methods). The LGN population atlas was carefully validated (SI Methods).

In a next step, the LGN population atlases were thresholded to 50% anatomical overlap across participants, and subsequently intersected with the registered single-subject qT_1_ maps (in template space) to extract the underlying qT_1_ values of the left and right LGN for each participant. We used the thresholded LGN population atlases rather than the participant-specific LGN masks to extract the underlying LGN qT_1_ distributions to ensure that the subsequent subdivision analyses were based on the same LGN voxels across participants.

*In-vivo* LGN subdivisions were identified by means of individual Gaussian mixture model fits as for the identification of *post-mortem* LGN subdivisions. Prior to Gaussian mixture model fitting, all of the extracted LGN qT_1_ distributions were normalized per hemisphere. The corrected qT_1_ distributions were centered on the same mean qT_1_ with equivalent standard deviation within each hemisphere across participants. The histogram normalization step ensured that model fits were based on a similar dynamic range of LGN qT_1_ data. Next, a Gaussian mixture model with two components, each representing one of the LGNs main subdivisions, was fit to each participant’s corrected left- and right-hemispheric qT_1_ distribution using the ‘curve_fit’ function in SciPy (44). The following anatomically motivated boundary conditions were employed on the Gaussian mixture models: (i) the volume of the Gaussian P-component is at least 50% of the total T_1_ distribution, which relates to prior anatomical findings in human *post-mortem* studies that the P subdivision occupies a larger part of total LGN volume than the M subdivision (12); (ii) the Gaussian P-component and the Gaussian M-component are centered between the 10^th^ and 50^th^ percentile and between the 50^th^ and 90^th^ percentile of the total qT_1_ distribution, respectively. This boundary condition relates to our findings from the *post-mortem* LGN subdivisions of shorter T_1_ relaxation in the P than M subdivision (see Results). (iii) Finally, both Gaussian M- and P-components were limited to standard deviations between 30% and 90% of the standard deviation of the total distribution. This ensured that not a single component alone (i.e., M or P) could span the entire qT_1_ data range. This boundary condition was motivated by the known two main subdivisions of the LGN (12), and our finding that T_1_ relaxation differs in subsections of the LGN that resemble these subdivisions in the *post-mortem* LGN data (see Results). The choice of range was motivated by the *post-mortem* results and was set to be centered on the percentage standard deviations of the identified *post-mortem* Gaussian M and P components.

All of the left and right-hemispheric Gaussian mixture model fits were subsequently subjected to the following quality control criteria: model fits were deemed anatomically implausible and were thus excluded when they returned a Gaussian mixture model wherein (i) not a single voxel had a higher probability for M than for P; and (ii) the number of M-classified voxels was larger than the number of P-classified voxels. 23 left-hemispheric and 24 right-hemispheric Gaussian mixture model fits out of 27 model fits per hemisphere fulfilled these criteria. Only for these model fits, each of the two individual Gaussian fits (i.e., M and P components) were normalized by the envelope of the Gaussian mixture model (i.e., whole distribution of M+P components) in the respective hemisphere to compute voxel-wise bilateral M and P distribution probability maps. This was done using the same procedure as for the identification of *post-mortem* LGN subdivisions. In a final step, the M and P distribution probability maps were combined for each hemisphere across participants to create bilateral population atlases of LGN M and P subdivisions.

### Data availability

Code of the employed in-house multi-contrast steady-state *post-mortem* qMRI fitting procedure as well as the *in-vivo* qMRI bilateral LGN and M/P subdivision population maps (in template and MNI 1mm standard space) are available for download at the Open Science Framework: https://doi.org/10.17605/OSF.IO/TQAYF.

## Results

### *Post-Mortem* MRI and Histology

The qT_1_ map of the *post-mortem* tissue sample revealed a clear contrast in qT_1_ between the LGN and surrounding gray and white matter structures (Fig. 1A). LGN segmentations of the two raters on the qT_1_ map were in excellent agreement (Dice coefficient 0.95), and resulted in an LGN volume of 87.6 mm^3^. Visual inspection of the qT_1_ map also revealed a clear qT_1_ contrast *within* the LGN: longer T_1_ relaxation seemed to coincide with the expected anatomical location of a ventral M subdivision, while shorter T_1_ relaxation seemed to coincide with a dorsal P subdivision (Fig. 1B & C). Fitting a Gaussian mixture model with two components, each representing one of the two main subdivisions of the LGN, to the underlying distribution of qT_1_ values in the LGN mask revealed one component centered around shorter qT_1_ values, and another component centered around longer qT_1_ values (Fig. 1D). A subsequent normalization of these two individual Gaussian components by the envelope of the Gaussian mixture model revealed a cluster of shorter T_1_ relaxation with high distribution probabilities in dorsal parts of the LGN (Fig. 1E), and a cluster of longer T_1_ relaxation with high distribution probabilities confined to ventral parts of the nucleus (Fig. 1F). Both clusters coincided with the expected anatomical location of LGN P (shorter T_1_ relaxation) and M (longer T_1_ relaxation) subdivisions, respectively. The identified dorsal and ventral subdivision contributed 75% and 25% to total LGN volume, respectively. This result is well in line with prior histological evidence on subdivisional size distributions in the human LGN of 72-81% for the parvocellular subdivision and 19-28% for the magnocellular subdivision (12).

**Fig. 1.**
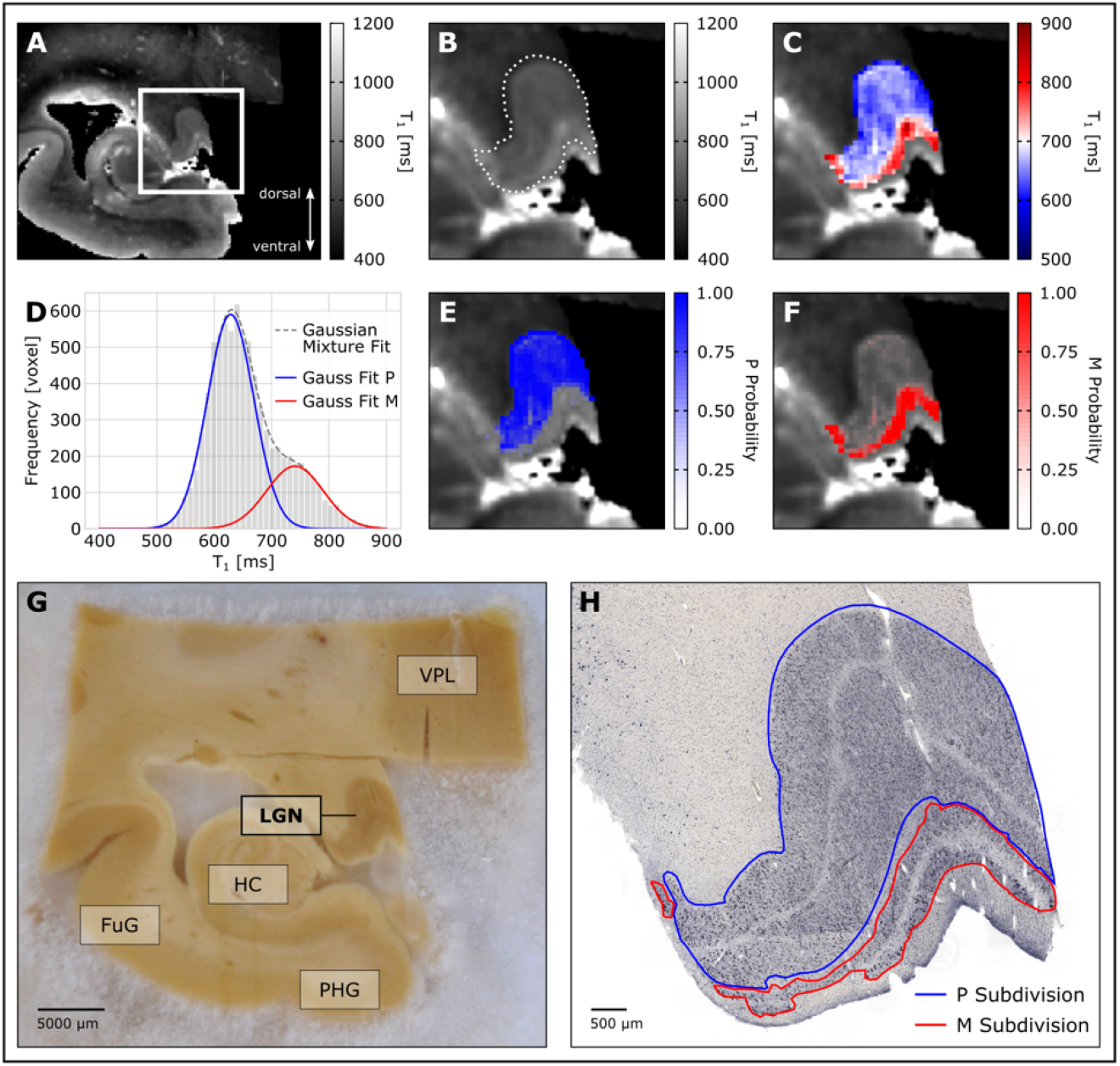
Identification of LGN subdivisions on ultra-high resolution qMRI and validation through microstructural histology in the same left-hemispheric tissue sample. **(A)** Coronal slice of the *post-mortem* qT_1_ map of the tissue sample. The white rectangle marks the location of the LGN. **(B)** Zoomed view of the LGN on the qT_1_ map. The dotted white line indicates the outline of the LGN. **(C)** Zoomed view of the LGN with an adapted color map to reflect the dynamic range of qT_1_ values within the LGN. Shorter qT_1_ values closely coincide with the anatomical location of a dorsal P subdivision, whereas longer qT_1_ values were found in the ventral M subdivision. **(D)** A two-component Gaussian mixture model was fit to the LGN qT_1_ distribution. The gray bars show the LGN qT_1_ histogram, outlined with the obtained Gaussian mixture model fit, as indicated by the gray dashed line. The blue and red curves correspond to the two components in the distribution of qT_1_ values, that were identified by the Gaussian mixture model. **(E)** Gaussian P-component, normalized by the envelope of the Gaussian mixture model reveals a cluster of short T_1_ relaxation with high distribution probabilities in dorsal parts of the LGN. **(F)** Gaussian M-component, normalized by the envelope of the Gaussian mixture model reveals a cluster of short T_1_ relaxation with high distribution probabilities in ventral parts of the nucleus. **(G)** Coronal view of the LGN tissue sample, showing the approximate same slice as shown in A-C & E-F. Anatomical labels are provided for spatial orientation. LGN = lateral geniculate nucleus, FuG = fusiform gyrus, HC = hippocampus, PHG = parahippocampal gyrus, VPL = ventral posterior lateral thalamic nucleus and other thalamic nuclei. **(H)** LGN slice, stained for anti-human neuronal protein C/D for neurons, shows characteristic M and P layering within the LGN. The figure shows the same slice as in G and the approximate same slice as in A-C & E-F. A typical four-layer LGN segment, consisting of two ventral M layers and two fused dorsal P layers, as often present in posterior parts of the nucleus, is visible (8). M and P subdivisions are shown to facilitate visual comparison with qMRI-based subdivision maps. The blue outline marks the P subdivision of the LGN. The red outline marks the M subdivision of the LGN.

Next, we compared our qMRI-based LGN subdivision maps against ground truth microstructural histology in the same LGN tissue sample. Immunohistochemical staining for neurons (anti-HuC/D) of the LGN sample (Fig. 1G&H) revealed a striking resemblence between histologically defined ventral M and dorsal P subdivisions and qT_1_-based mappings of LGN subdivisions (Fig. 1E & F).

A layer-specific stereological analysis of the neuronal cell density within the LGN (Fig. 2A & B) revealed as expected (7, 12) a lower cell density in M than P layers. A layer-specific optical density analysis of immunohistochemical staining for myelin (anti-MBP) revealed a stronger myelination of the P layers, compared to the M layers in the LGN (Fig. 2C & D). A layer-specific optical density analysis of histochemical staining for ferric iron (Perls’ Prussian blue) yielded no clear distinction in terms of iron content between LGN M and P subdivisons; but interestingly revealed greater iron deposits in the parvocellular segment of fused layers P4/6, which receive visual input from the contralateral eye (Fig. 2E & F).

**Fig. 2.**
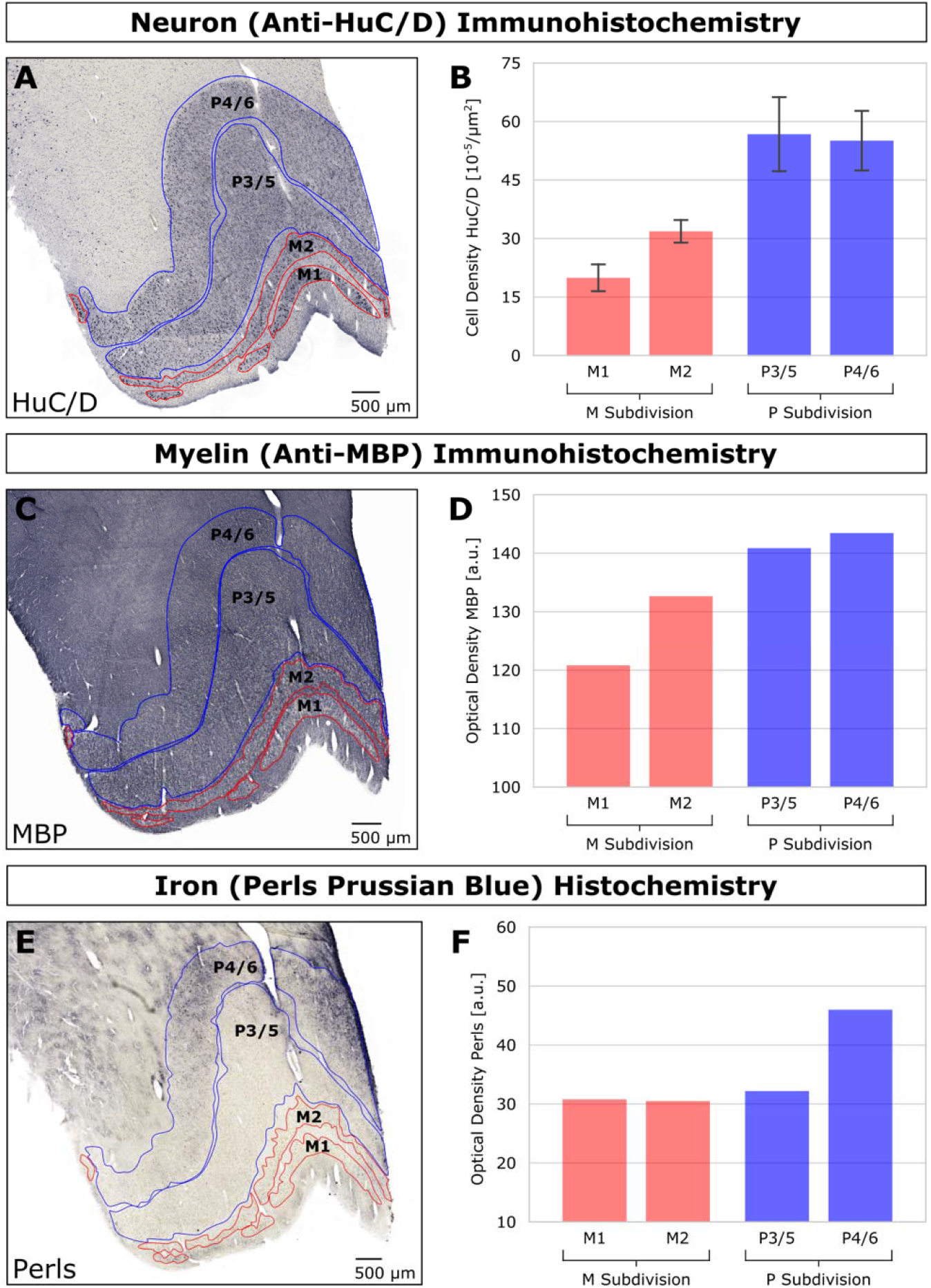
Histological assessments of microstructural tissue differences between LGN layers. **(A)** Immunohistochemical staining for neurons (anti-HuC/D) of the LGN specimen. Individual LGN P layers are outlined in blue. LGN M layers are outlined in red. The two layers P4/6 (contralateral input) and P3/5 (ipsilateral input) are fused in this posterior slice of the LGN (same slice as in Fig. 1H). **(B)** Bar graph of stereological assessment of the estimated neuronal cell density for individual P (blue) and M (red) layers from data shown in A. Error bars indicate ± 1 SD. **(C)** Immunohistochemical staining for myelin (anti-MPB) of the LGN specimen. LGN P layers are outlined in blue. LGN M layers are outlined in red. The figure shows the approximate same slice as in Fig. 1H and Fig. 2A. **(D)** Bar graph of the MBP optical density of individual P (blue) and M (red) layers from data shown in C. In contrast to the cell density measurements shown in (B), the MBP optical density measurements yielded exactly one value per layer. Therefore, error bars do not apply (SI Methods). **(E)** Histochemical staining for ferric iron (Perls’ Prussian blue, PB) of the LGN specimen. Individual LGN P layers are outlined in blue. LGN M layers are outlined in red. The figure shows approximately the same slice as in Fig. 1H and Fig. 2A & B. **(F)** Bar graph of the PB optical density of individual P (blue) and M (red) layers from data shown in A. The PB optical density measurements yielded exactly one value per layer. Therefore, error bars do not apply (SI Methods).

Taken together, the histological findings suggest that the higher cell density in P layers of the LGN increases the net myelin density of the P subdivision, which could explain the observation of overall shorter T_1_ relaxation of the P subdivision than M subdivision in the qT_1_ map.

### *In-vivo* MRI

Inter-rater reliability measures of the LGN segmentations on the N=27 *in-vivo* qT_1_ maps indicated high agreement between the two raters (mean Dice coefficients: left LGN=0.88±0.02; right LGN=0.89±0.02). The segmentation procedure resulted in mean LGN volumes of 113.5±13.3mm^3^ and 120.9±14.0mm^3^ for the left and right LGN, respectively. LGN volumes were significantly correlated across hemispheres (*R*=0.73, p=2.2×10^−5^), and were significantly greater in the right than in the left hemisphere (*t*(26)=3.9, p=0.7×10^−3^). Underlying mean qT_1_ values of the left and right LGN masks showed a high correspondence (left LGN: 1469.9±61.2ms, right LGN: 1469.2±60.9ms; *R*=0.93, p=3.8×10^−12^), and did not significantly differ from each other (*t*(26)=0.16, p=0.88). A normalization of the LGN masks to the study-specific qT_1_ group template (Fig. 3A and Fig. S1C), followed by overlaying of the masks for each hemisphere, resulted in bilateral LGN population atlases that showed a strong correspondence across participants with the underlying anatomy (Fig. 3A & B). The bilateral LGN population atlas, thresholded to 50% anatomical overlap across participants (Fig. 4A & B), was subsequently employed to extract the individual LGN qT_1_ distributions of all participants in each hemisphere. For the identification of *in-vivo* LGN subdivisions, two-component Gaussian mixture models were fit to the extracted single-subject LGN qT_1_ distributions, and the resulting segmentation of the subdivisions were combined across participants for each hemisphere. The resulting population maps of the P and M subdivisions revealed a similar pattern as the *post-mortem* qMRI results: P-classified voxels with shorter T_1_ relaxation showed the largest anatomical overlap across participants in dorsal parts of the LGN (Fig. 4C & D), whereas M-classified voxels with longer T_1_ relaxation showed the largest anatomical overlap in ventral parts of the nucleus (Fig. 4E & F). Thresholding of these P and M subdivision populations maps to 50% anatomical overlap across participants revealed two spatially distinct, non-overlapping clusters, which in turn coincided with the expected anatomical location of a dorsal P and a ventral M LGN subdivision (Fig. 4G & H). The ratio of the sums of the identified P and M components across participants resulted in average contributions of 68.1±14.8% (P-component) versus 31.9±14.8% (M-component) to total LGN volume in the left hemisphere; and 66.0±15.3% (P-component) versus 34.0±15.3% (M-component) in the right hemisphere.

**Fig. 3.**
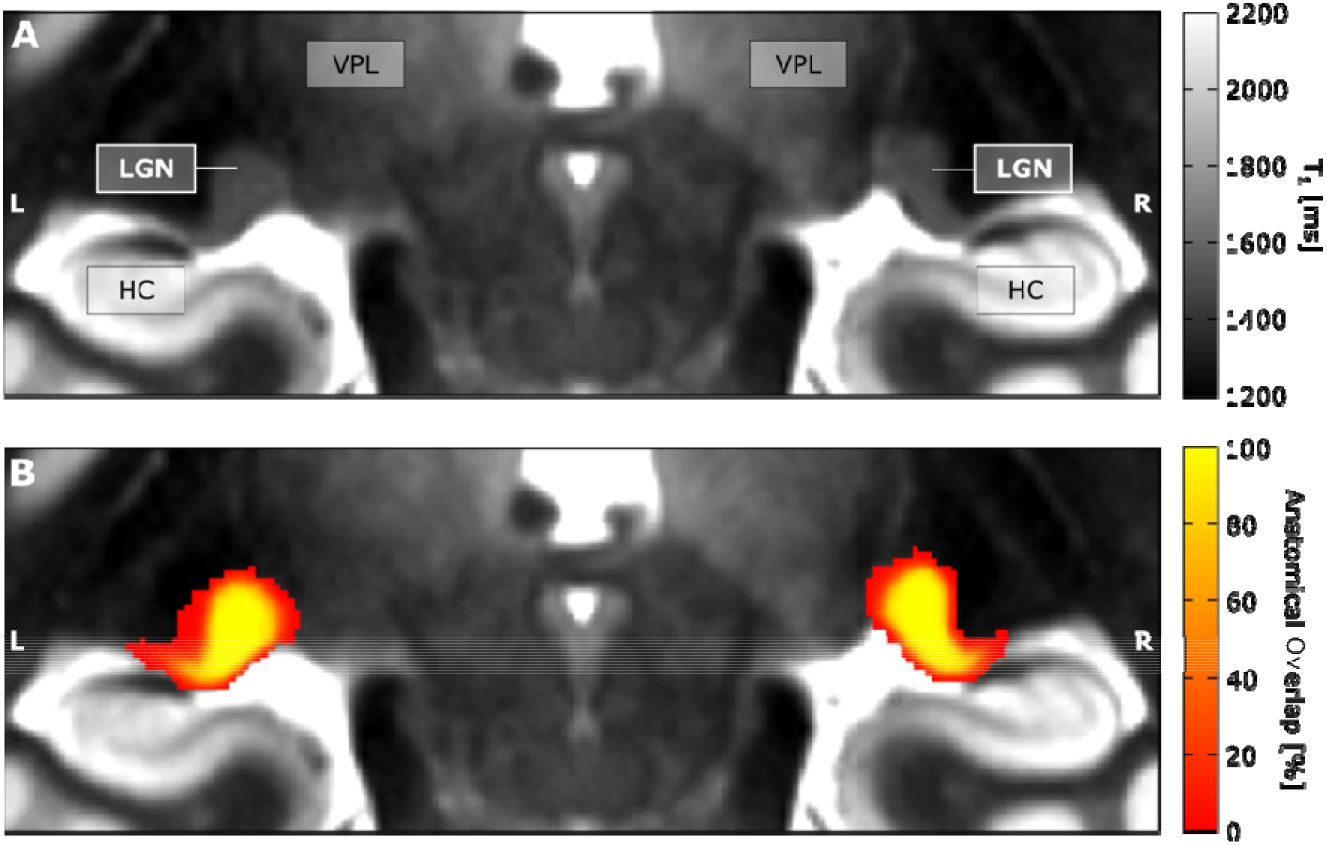
Coronal view of the study-specific qT_1_ group template along with the bilateral *in-vivo* LGN population atlas. **(A)** The figure shows a coronal slice of the study-specific qT_1_ group template, centered on the lateral geniculate nuclei. Anatomical labels are provided for spatial orientation and comparison to Fig. 1G: LGN = lateral geniculate nucleus, HC = hippocampus, VPL = ventral posterior lateral thalamic nucleus and other thalamic nuclei. **(B)** Study-specific qT_1_ group template as shown in A, overlaid with the left and right LGN population atlas. Color coding indicates the anatomical overlap in LGN location across the N=27 participants.

**Fig. 4.**
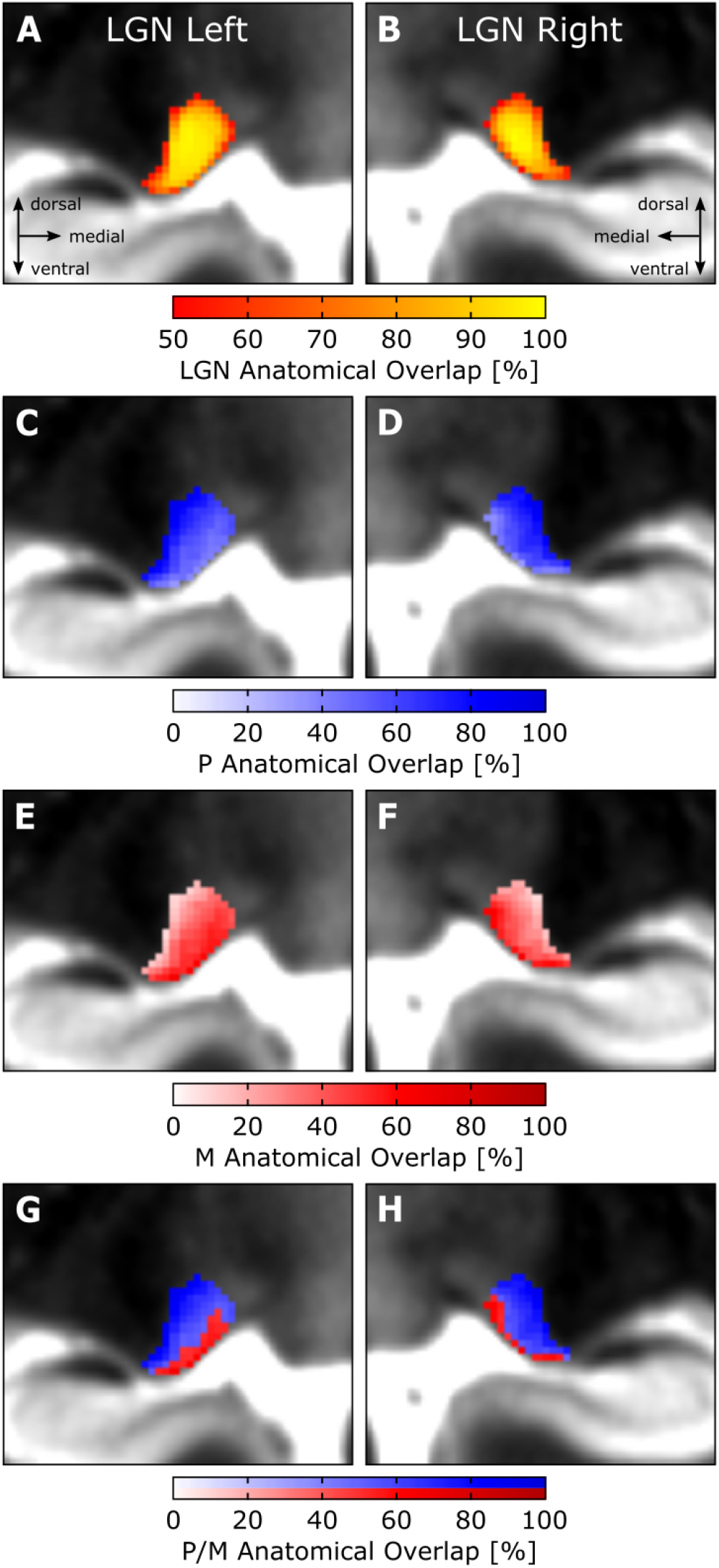
*In-vivo* LGN subdivisions overlaid on a slice of the study-specific qT_1_ group template in coronal view. **(A-B)** Zoomed view of the left (A) and right (B) LGN population atlas based on N=27 participants, thresholded to 50% anatomical overlap across participants. **(C-D)** Zoomed view of the population atlas of the left (C) and right (D) LGN P subdivision. Color coding indicates the anatomical overlap in P-classified voxels across participants. **(E-F)** Zoomed view of the population atlas of the left (E) and right (F) LGN M subdivision. Color coding indicates the anatomical overlap in M-classified voxels across participants. **(G-H)** Zoomed view of the left (G) and right (H) LGN P and M subdivision population maps, thresholded to 50% anatomical overlap across participants. The blue-shaded part of the color bar indicates the anatomical overlap for the dorsal P cluster, while the red-shaded part of the color bar indicates the anatomical overlap for the ventral M cluster. The P and M subdivision population maps in C-H are based on N=23 and N=24 participants in the left and right hemisphere, respectively (see Methods).

## Discussion

Recent advances in quantitative MRI (qMRI) enable researchers to gain insights into the microstructural tissue properties of the human brain (24). In the present study, we utilized these technological advances for mapping the human LGN, a key structure in the early visual pathway (1). We demonstrated that a differentiation of the LGN and its two functionally distinct main subdivisions is possible based on microstructure-informed qMRI contrasts alone. To this end, we employed ultra-high resolution qMRI of a *post-mortem* LGN specimen and high-resolution *in-vivo* qMRI in a large sample of N=27 participants. Through histological examination, we showed how the observed qMRI contrast relates to differences in cell and myelin density between LGN subdivisions. The study overcomes long-standing technical challenges to reveal LGN subdivisions in humans *in-vivo*. It paves the way for imaging subsections of the LGN to obtain a better understanding of its function and microstructure in health and disease.

### Microstructural properties that enable a differentiation of M/P subdivisions

The quantitative assessment of the LGN and a differentiation of its main subdivisions was possible based on the longitudinal relaxation time T_1_. The observed qT_1_ contrast directly related to cyto-and myeloarchitectonic differences between LGN M and P layers. A histological examination of the *post-mortem* LGN sample confirmed a higher cell density in P layers compared to the M layers of the LGN (7, 12, 31, 32). LGN P layers were further found to have a higher myelin density than M layers. In contrast to cell and myelin density, a comparison of iron content did not yield a clear differentiation between M and P layers. The finding of a higher myelin density in P than M layers seems at a first glance counter-intuitive, as M axons are known to be more myelinated than P axons (32–36). Our findings suggest that the greater cell density in P layers of the LGN acts as mediating factor for the observed differences in myelin density between P and M layers: the higher density of less myelinated P axons increases the net myelin density of the P layers of the LGN, as compared to a smaller amount of more heavily myelinated M axons. Given the inverse relationship between myelin density per voxel and T_1_ relaxation (25, 27), this would explain the observed *post-mortem* qT_1_ contrast of shorter T_1_ relaxation in P layers and longer T_1_ relaxation in M layers of the LGN. The mediating role of the cell density highlights the crucial need to take cytoarchitectonic tissue features, such as the cell density, into account when making inferences about myelin density from T_1_ relaxometry. The possibility of dissociating LGN subdivisions based on their microstructural tissue properties in *post-mortem* qMRI acquisitions is intriguing as it poses a valuable opportunity to also assess these subdivisions in humans *in-vivo*. In line with this argumentation, we have shown that microstructure-informed qMRI can guide the differentiation of LGN subdivisions *in-vivo*: similar to the *post-mortem* results, two components of shorter and longer T_1_ relaxation were identified, that coincided with the anatomical location of dorsal P and ventral M subdivisions, respectively.

### Large inter-individual differences

The observed relative size contributions of the identified *post-mortem* and *in-vivo* LGN subdivisions to total LGN volume were well in line with prior histological assessments in the human LGN: 72-81% for the parvocellular subdivision and 19-28% for the magnocellular subdivision (12). The identified *in-vivo* LGN subdivisions, however, showed a large degree of variation across participants. This variation might be due to genuine inter-individual differences: from *post-mortem* studies we know that the human LGN has an approximately twofold inter-individual variability in volume (12). This large variation in LGN volume across participants could also extend to large inter-individual differences in M and P subdivision volumes and thus explain the large inter-individual differences in subdivision volume found in the present study. However, there is also the possibility that the amount of inter-individual variability could be reduced by further technological improvements. First, there is an inherent trade-off between brain coverage, resolution and sensitivity in MRI acquisitions (54). The image resolution constraints arising from whole-brain acquisitions might have hampered *in-vivo* mappings of LGN subdivisions. Custom-tailored *in-vivo* qMRI acquisitions of the LGN with reduced coverage of only a part of the brain for the benefit of increased image resolution are likely to enhance subdivisional LGN contrast. Second, we showed that a single qMRI contrast (i.e., qT_1_) can reveal subdivisions also *in-vivo*. However, using multi-modal qMRI approaches, such as multi-parametric mapping (45), might further boost the sensitivity of microstructural qMRI contrast to LGN subdivisional mapping.

### Why is qMRI better than fMRI for assessing LGN subdivisions?

Conventionally, *in-vivo* MR imaging of the LGN and its subdivisions has faced severe technical challenges due to the LGNs small extent and deep location within the brain. A few recent studies have demonstrated that more M and P-like activation clusters in the LGN can be identified *in-vivo* using extensive fMRI protocols (19, 38, 39). The method proposed here of using qMRI to dissociate between LGN subdivisions offers significant advantages over fMRI acquisitions: (i) In contrast to fMRI, structural MRI is characterized by a higher signal-to-noise ratio and allows for acquisitions with higher spatial resolution; (ii) structural MRI acquisitions are generally less taxing for participants as no lengthy functional design, often consisting of multiple repetitions of experimental stimuli, is required; and (iii) qMRI allows to draw conclusions about the underlying microstructure, such as tissue myelination (24). The ability to assess microstructural information about the LGN and its subdivisions *in-vivo* offers novel opportunities to unravel their specific contribution to clinical conditions such as multiple sclerosis (20) and glaucoma (19). While 7 Tesla MRI systems are increasingly available in clinical environments, we expect that custom-tailored multi-modal qMRI acquisitions will also enable quantitative assessments of the LGN and its subdivisions even at lower field strengths.

### The role of iron

Besides the findings related to our main aim, we made three further interesting observations, two relating to iron content and one relating to the lateralization of the LGN. First, while we did not find a clear differentiation in iron content between M and P subdivisions, we did observe greater iron deposits in layers P4/6 (Fig. 2E & F), which receive visual input from the contralateral eye (8). This finding is intriguing, as it suggests a different composition of eye-specific contralateral and ipsilateral P layers. In histological preparations, contralateral and ipsilateral P layers show a relatively uniform morphological appearance (8, 12), (Fig. 1H & Fig. 2A). However, functional differences between contralateral and ipsilateral P layers with respect to response latencies of their constituent neurons have been reported previously in non-human primate research (55). Whether these functional differences also apply to humans and whether they relate to local differences in iron content is currently unknown. To our knowledge there is no study that has yet examined layer-specific differences in iron content in the human LGN. Multiple studies have shown that iron concentrations in the human brain accumulate with aging (56, 57). As our LGN sample was obtained from a patient of advanced age (89yrs), we cannot exclude the possibility that the elevated iron deposits in layers P4/6 are due to age-related changes in brain iron concentration levels. However, the specificity of elevated iron deposits in contralateral layers P4/6, and the otherwise low iron content in the thalamus (27, 58), make this explanation unlikely.

Second, in light of the elevated iron deposits in LGN layers P4/6, and the inverse relationship between iron content and T_1_ relaxation (27), one could have expected an additional qT_1_ component centered around the shorter qT_1_ range of the identified P component in the *post-mortem* qT_1_ LGN distribution. Nonetheless, the *post-mortem* qT_1_ LGN distribution bore no sign of an additional qT_1_ component that captures the elevated iron deposits in contralateral layers P4/6 (Fig. 1A-F). The lack of such an iron-specific qT_1_ component is consistent with the concept that iron content, as compared to myelin density, only has a relatively small contribution to T_1_ relaxation (27).

### Lateralization of LGN volume

The third interesting finding concerned the hemispheric differences in LGN volumes in our *in-vivo* qMRI data. LGN volumes were significantly greater in the right than in the left hemisphere. Indications for greater right-hemispheric LGN volumes in healthy control participants have been reported before in MRI (59, 60) and histological studies (12). Other MRI studies did not find evidence for such lateralization effects (2, 61, 62). One explanation for these variable findings across studies might relate to the large inter-individual variability in LGN volume in humans (12); and that previous studies with low or modest sample sizes lack sufficient statistical power to reliably detect potential inter-hemispheric differences in LGN volume (2, 12).

### Limitations

The human LGN is a layered structure consisting of not only parvo-and magnocellular layers, but also koniocellular layers (12). Thus far, it has only been possible to dissociate the M and P subdivisions of the LGN using MRI. K neurons are the smallest among the three neuron types, which form narrow intercalated layers in the LGN (63). Because of their small size, mapping of these layers via MRI has to date not been possible due to limited image resolution. The functional role of K neurons is largely unexplored (63). However, they have been implicated in the direct V1-bypassing connection between the LGN and visual motion area MT in non-human primate research (64–66) – a pathway that has also been linked to specific clinical conditions in humans such as blindsight (67, 68) and developmental dyslexia (2, 68).

### Implications

Only recent advances in MRI have made it possible to investigate the LGN and its subdivisions in humans *in-vivo*. Therefore to-date we are only starting to understand the LGNs complex role for human perception and cognition in health (1) and disease (2–5). We introduced here a microstructure-informed strategy to map the LGN and its subdivisions both *post-mortem* and *in-vivo* using qMRI – a strategy, which offers potential applications also for sub-parcellations of other subcortical nuclei with distinctive cyto-and myeloarchitectonic tissue differences. By using two orthogonal qMRI acquisition schemes, we show qT_1_ differences between LGN M and P subdivisions, which are directly related to cyto- and myeloarchitectonic differences between subdivisions, as confirmed through histological validation. The method proposed here paves the way for a detailed assessment of LGN function and microstructure in humans at an unprecedented level of image resolution that is more readily comparable to data from animal research. It also provides a novel opportunity for investigating the contribution of selective impairments in LGN subdivisions to clinical disorders such as multiple sclerosis (20), glaucoma (19), and developmental dyslexia (2, 3). With high-field MRI systems being more readily available, we are confident that the qMRI contrast demonstrated here will constitute an important future milestone for assessing LGN function and dysfunction in humans both in neuroscientific and clinical settings. As a contribution to this endeavor, we have made the *in-vivo* LGN population atlas and M and P subdivision population maps presented here publicly available to facilitate standardized future studies on the human LGN.

## References

1. Y. B. Saalmann, S. Kastner, Cognitive and perceptual functions of the visual thalamus. Neuron 71, 209–23 (2011).

2. C. Müller-Axt, A. Anwander, K. von Kriegstein, Altered structural connectivity of the left visual thalamus in developmental dyslexia. Curr. Biol. 27, 3692–3698 (2017).

3. M. S. Livingstone, G. D. Rosen, F. W. Drislane, A. M. Galaburda, Physiological and anatomical evidence for a magnocellular defect in developmental dyslexia. Proc. Natl. Acad. Sci. U.S.A. 88, 7943–7947 (1991).

4. K. A. Dorph-Petersen, et al., Volume and neuron number of the lateral geniculate nucleus in schizophrenia and mood disorders. Acta Neuropathol. 117, 369–384 (2009).

5. Y. H. Yücel, Q. Zhang, R. N. Weinreb, P. L. Kaufman, N. Gupta, Effects of retinal ganglion cell loss on magno-, parvo-, koniocellular pathways in the lateral geniculate nucleus and visual cortex in glaucoma. Prog. Retin. Eye Res. 22, 465–481 (2003).

6. B. U. Forstmann, G. de Hollander, L. van Maanen, A. Alkemade, M. C. Keuken, Towards a mechanistic understanding of the human subcortex. Nat. Rev. Neurosci. 18, 57–65 (2016).

7. J. J. Nassi, E. M. Callaway, Parallel processing strategies of the primate visual system. Nat. Rev. Neurosci. 10, 360–372 (2009).

8. T. L. Hickey, R. W. Guillery, Variability of laminar patterns in the human lateral geniculate nucleus. J. Comp. Neurol. 183, 221–246 (1979).

9. D. H. O’Connor, M. M. Fukui, M. A. Pinsk, S. Kastner, Attention modulates responses in the human lateral geniculate nucleus. Nat. Neurosci. 5, 1203–1209 (2002).

10. P. G. Mihai, et al., Modulation of tonotopic ventral medial geniculate body is behaviorally relevant for speech recognition. eLife 8, 1–28 (2019).

11. Y. B. Saalmann, S. Kastner, Gain control in the visual thalamus during perception and cognition. Curr. Opin. Neurobiol. 19, 408–414 (2009).

12. T. J. Andrews, S. D. Halpern, D. Purves, Correlated size variations in human visual cortex, lateral geniculate nucleus, and optic tract. J. Neurosci. 17, 2859–2868 (1997).

13. A. Weibull, H. Gustavsson, S. Mattsson, J. Svensson, Investigation of spatial resolution, partial volume effects and smoothing in functional MRI using artificial 3D time series. NeuroImage 41, 346–353 (2008).

14. M. Livingstone, D. Hubel, Segregation of form, color, movement, and depth: anatomy, physiology, and perception. Science 240, 740–749 (1988).

15. K. A. Schneider, S. Kastner, Effects of sustained spatial attention in the human lateral geniculate nucleus and superior colliculus. J. Neurosci. 29, 1784–1795 (2009).

16. R. N. Denison, M. A. Silver, Distinct contributions of the magnocellular and parvocellular visual streams to perceptual selection. J. Cogn. Neurosci. 24, 246–259 (2012).

17. E. M. Callaway, Structure and function of parallel pathways in the primate early visual system. J. Physiol. 566, 13–19 (2005).

18. N. Gupta, L. C. Ang, L. N. de Tilly, L. Bidaisee, Y. H. Yücel, Human glaucoma and neural degeneration in intracranial optic nerve, lateral geniculate nucleus, and visual cortex. Br. J. Ophthalmol. 90, 674–678 (2006).

19. P. Zhang, W. Wen, X. Sun, S. He, Selective reduction of fMRI responses to transient achromatic stimuli in the magnocellular layers of the LGN and the superficial layer of the SC of early glaucoma patients. Hum. Brain Mapp. 37, 558–569 (2016).

20. N. Evangelou, et al., Size-selective neuronal changes in the anterior optic pathways suggest a differential susceptibility to injury in multiple sclerosis. Brain 124, 1813–1820 (2001).

21. M. J. Thurtell, et al., Evaluation of optic neuropathy in multiple sclerosis using low-contrast visual evoked potentials. Neurology 73, 1849–1857 (2009).

22. J. Stein, To see but not to read; the magnocellular theory of dyslexia. Trends Neurosci. 20, 147–152 (1997).

23. E. Milne, et al., High motion coherence thresholds in children with autism. J. Child Psychol. Psychiatry 43, 255–263 (2002).

24. W. van der Zwaag, A. Schäfer, J. P. Marques, R. Turner, R. Trampel, Recent applications of UHF-MRI in the study of human brain function and structure: a review. NMR Biomed. 29, 1274–1288 (2016).

25. S. Geyer, M. Weiss, K. Reimann, G. Lohmann, R. Turner, Microstructural parcellation of the human cerebral cortex - from Brodmann’s post-mortem map to in vivo mapping with high-field magnetic resonance imaging. Front. Hum. Neurosci. 5, 1–7 (2011).

26. A. Lutti, F. Dick, M. I. Sereno, N. Weiskopf, Using high-resolution quantitative mapping of R1 as an index of cortical myelination. NeuroImage 93, 176–188 (2014).

27. C. Stüber, et al., Myelin and iron concentration in the human brain: a quantitative study of MRI contrast. NeuroImage 93, 95–106 (2014).

28. P. L. Bazin, et al., A computational framework for ultra-high resolution cortical segmentation at 7 Tesla. NeuroImage 93, 201–209 (2014).

29. M. D. Waehnert, et al., A subject-specific framework for in vivo myeloarchitectonic analysis using high resolution quantitative MRI. NeuroImage 125, 94–107 (2016).

30. E. Kuehn, et al., Body topography parcellates human sensory and motor cortex. Cereb. Cortex 27, 3790–3805 (2017).

31. B. Dreher, Y. Fukada, R. W. Rodieck, Identification, classification and anatomical segregation of cells with X-like and Y-like properties in the lateral geniculate nucleus of old-world primates. J. Physiol. 258, 433–452 (1976).

32. R. Hassler, “Comparative anatomy of the central visual systems in day-and night-active primates” in Evolution of the Forebrain, (Springer, 1966), pp. 419–434.

33. W. H. Merigan, J. H. R. Maunsell, How parallel are the primate visual pathways? Annu. Rev. Neurosci. 16, 369–402 (1993).

34. J. Stein, “The neurobiology of reading difficulties” in Basic Functions of Language, Reading and Reading Disability, Neuropsychology and Cognition., E. Witruk, A. D. Friederici, T. Lachmann, Eds. (Springer US, 2002), pp. 199–211.

35. A. A. Beaton, “The magnocellular deficit hypothesis” in Dyslexia, Reading and the Brain: A Sourcebook of Psychological and Biological Research, 1st Ed., (Psychology Press, 2004), pp. 262–281.

36. A. Yoonessi, A. Yoonessi, Functional assessment of magno, parvo and konio-cellular pathways; current state and future clinical applications. J. Ophthalmic Vis. Res. 6, 119–126 (2011).

37. S. M. Sherman, J. R. Wilson, J. H. Kaas, X- and Y-cells in the dorsal lateral geniculate nucleus of the owl monkey (Aotus trivirgatus). Science 192, 475–477 (1976).

38. R. N. Denison, A. T. Vu, E. Yacoub, D. A. Feinberg, M. A. Silver, Functional mapping of the magnocellular and parvocellular subdivisions of human LGN. NeuroImage 102, 358–369 (2014).

39. P. Zhang, H. Zhou, W. Wen, S. He, Layer-specific response properties of the human lateral geniculate nucleus and superior colliculus. NeuroImage 111, 159–166 (2015).

40. M. A. Bernstein, K. F. King, X. J. Zhou, Handbook of MRI Pulse Sequences (Elsevier Academic Press, 2004).

41. J. E. Iglesias, et al., Effect of Fluorinert on the histological properties of Formalin-fixed human brain tissue. J. Neuropathol. Exp. Neurol. 12, 1085–1090 (2018).

42. G. Helms, H. Dathe, P. Dechent, Quantitative FLASH MRI at 3T using a rational approximation of the Ernst equation. Magn. Reson. Med. 59, 667–672 (2008).

43. R. Müller, T. Lenich, E. Kirilina, H. E. Möller, Application of an RF current mirror for MRI transmit coils in ISMRM 27th Annual Meeting & Exhibition, (2019).

44. P. Virtanen, et al., SciPy 1.0: fundamental algorithms for scientific computing in Python. Nat. Methods 17, 261–272 (2020).

45. N. Weiskopf, M. F. Callaghan, O. Josephs, A. Lutti, S. Mohammadi, Estimating the apparent transverse relaxation time (R2*) from images with different contrasts (ESTATICS) reduces motion artifacts. Front. Neurosci. 8, 1–10 (2014).

46. N. J. Tustison, et al., N4ITK: Improved N3 bias correction. IEEE Trans. Med. Imag. 29, 1310–1320 (2010).

47. J. P. Marques, R. Gruetter, New developments and applications of the MP2RAGE sequence-focusing the contrast and high spatial resolution R1 mapping. PLoS One 8, 1–11 (2013).

48. J. P. Marques, et al., MP2RAGE, a self bias-field corrected sequence for improved segmentation and T1-mapping at high field. NeuroImage 49, 1271–1281 (2010).

49. L. R. Dice, Measures of the amount of ecologic association between species. Ecology 26, 297–302 (1945).

50. C. L. Tardif, et al., Open Science CBS Neuroimaging Repository: sharing ultra-high-field MR images of the brain. NeuroImage 124, 1143–1148 (2016).

51. R. C. Oldfield, The assessment and analysis of handedness: the Edinburgh inventory. Neuropsychologia 9, 97–113 (1971).

52. B. B. Avants, C. L. Epstein, M. Grossman, J. C. Gee, Symmetric diffeomorphic image registration with cross-correlation: Evaluating automated labeling of elderly and neurodegenerative brain. Med. Image Anal. 12, 26–41 (2008).

53. B. B. Avants, et al., The optimal template effect in hippocampus studies of diseased populations. NeuroImage 49, 2457–2466 (2010).

54. L. Huber, et al., Layer-dependent functional connectivity methods. Prog. Neurobiol., 101835 (2020).

55. J. H. R. Maunsell, et al., Visual response latencies of magnocellular and parvocellular LGN neurons in macaque monkeys. Vis. Neurosci. 16, 1–14 (1999).

56. P. Ramos, et al., Iron levels in the human brain: A post-mortem study of anatomical region differences and age-related changes. J. Trace Elem. Med. Biol. 28, 13–17 (2014).

57. R. J. Ward, F. A. Zucca, J. H. Duyn, R. R. Crichton, L. Zecca, The role of iron in brain ageing and neurodegenerative disorders. Lancet Neurol. 13, 1045–1060 (2014).

58. W. D. Rooney, et al., Magnetic field and tissue dependencies of human brain longitudinal 1H2O relaxation in vivo. Magn. Reson. Med. 57, 308–318 (2007).

59. M. Li, et al., Quantification of the human lateral geniculate nucleus in vivo using MR imaging based on morphometry: Volume loss with age. Am. J. Neuroradiol. 33, 915–921 (2012).

60. A. Papadopoulou, et al., Damage of the lateral geniculate nucleus in MS: Assessing the missing node of the visual pathway. Neurology 92, 2240–2249 (2019).

61. L. Mcketton, K. R. Kelly, K. A. Schneider, Abnormal lateral geniculate nucleus and optic chiasm in human albinism. J. Comp. Neurol. 522, 2680–2687 (2014).

62. M. Giraldo-Chica, J. P. Hegarty II, K. A. Schneider, Morphological differences in the lateral geniculate nucleus associated with dyslexia. NeuroImage Clin. 7, 830–836 (2015).

63. S. H. Hendry, R. C. Reid, The koniocellular pathway in primate vision. Annu. Rev. Neurosci. 23, 127–153 (2000).

64. L. C. Sincich, K. F. Park, M. J. Wohlgemuth, J. C. Horton, Bypassing V1: A direct geniculate input to area MT. Nat. Neurosci. 7, 1123–1128 (2004).

65. C. E. Warner, Y. Goldshmit, J. A. Bourne, Retinal afferents synapse with relay cells targeting the middle temporal area in the pulvinar and lateral geniculate nuclei. Front. Neuroanat. 4, 1–16 (2010).

66. M. C. Schmid, et al., Blindsight depends on the lateral geniculate nucleus. Nature 466, 373–377 (2010).

67. H. Bridge, et al., Visual activation of extra-striate cortex in the absence of V1 activation. Neuropsychologia 48, 4148–4154 (2010).

68. S. Rima, M. Christoph Schmid, V1-bypassing thalamo-cortical visual circuits in blindsight and developmental dyslexia. Curr. Opin. Physiol. 16, 14–20 (2020).

