## Supplementary Information for "Mapping the Human Visual Thalamus and its Cytoarchitectonic Subdivisions Using Quantitative MRI"

DRAFT

### SI Materials and Methods

#### *Post-Mortem* MRI and Histology

Gaussian Mixture Model Components. To determine the adequate number of compartments (i.e., the number of Gaussian components) in the Gaussian mixture model, we used a data-driven approach to evaluate the fitting accuracy with respect to model complexity. To this end, the model fitting accuracy was related to the number of required parameters. The objective was to identify the most accurate model for the number of parameters involved.

Three different Gaussian mixture models with one (1C), two (2C), or three (3C) components were fit to the distribution of $qT_1$ values within the LGN of the ultra-high-resolution *post-mortem* MRI data. For all models, the LGN $qT_1$ distribution was calculated using 50 equidistant bins covering the entire data range, spread from $qT_{1,min}$=400ms to $qT_{1,max}$=900ms. The fitting accuracy was calculated utilizing the root mean square deviation (RMSD) between the histogram and model prediction. To include the model complexity in the assessment, the RMSD values were weighted by the number of model parameters of each Gaussian mixture model (i.e., the number of Gaussian components times three parameters [mean, standard deviation, amplitude] per component). The resulting weighted RMSDs (wRMSD) were $wRMSD_{1C}$=1496ms, $wRMSD_{2C}$=154ms, $wRMSD_{3C}$=203ms. Hence, the two component Gaussian mixture model achieved the lowest weighted error for its given model complexity and, therefore, was chosen to most accurately represent the $qT_1$ variation within the LGN.

HuC/D and MBP Immunohistochemistry. For histological examination, the LGN tissue block was cryoprotected in 30% sucrose and cut into 30µm consecutive sections using a Jung Histoslide 2000 freezing microtome (Leica, Wetzlar, Germany) equipped with a Hyrax 30 freezing unit (Carl Zeiss, Jena, Germany). The LGN comprised 5.4mm (180 sections from posterior to anterior pole) in total. Every seventh 30µm section was used for a series of immunohistochemical stains to cover the LGN in slices compatible with the *post-mortem* MR acquisition resolution (immunohistochemistry: 7x30µm = 210µm; MR acquisition resolution = 220µm). After washing the free-floating slices in phosphate-buffered saline with Tween (PBS-Tween), slices were pre-treated for antigen retrieval following previously published procedures (1). Afterwards, another washing step was employed, and samples were incubated in blocking solution (2% bovine serum albumin (BSA), 0.3% milk powder and 0.5% donkey normal serum (DNS) in PBS) for 1h at room temperature to avoid unspecific binding of antibodies. Next, slices were incubated with primary antibodies, HuC/D (mouse; 1:500; A21271; ThermoFisher Scientific, Waltham, MA, USA) or MBP (rat; 1:400; NB600-717; Novus Biologicals, Littleton, CO, USA) for 48h at 4°C in blocking solution. Subsequently, sections were washed in PBS-Tween and incubated for 1h in biotinylated secondary antibody solution (donkey-anti-mouse or donkey-anti-rat; 1:1000; Dianova, Hamburg, Germany) containing PBS-Tween and blocking solution (1:2). Again, sections were washed in PBS-Tween and incubated in streptavidin (1:2000; Extravidin®; Sigma Aldrich, St. Louis, MO, USA). After washing in PBS-Tween and Tris-Hydrochloride (Tris-HCl; pH 8.0), samples were batch-wise developed in 3,3'-Diaminobenzidine (DAB; Sigma Aldrich) and nickel-ammonium sulphate (Sigma Aldrich) for 3min under visual control. Last washing steps were performed with Tris-HCl (pH 8.0) and

DRAFT

PBS before samples were mounted onto microscopic slides, air-dried and coverslipped with Entellan® (Merck, Darmstadt, Germany).

Perls' Prussian Blue Histochemistry. Cryo-cut 30µm samples were mounted onto microscopic slides and air-dried overnight. Sections were then washed in distilled and double-distilled water. Perls' Prussian blue solution consisted of 1:1 freshly mixed 5% potassium hexacyanoferrate-II and 5% hydrochloric acid. Samples were immersed and incubated at 37°C for 2h (2, 3). Samples were then washed in phosphate-buffered saline (PBS) and Tris-Hydrochloride (Tris-HCl; pH 8.0) and for intensification of stain developed in 3,3'-diaminobenzidine (DAB; Sigma Aldrich, St. Louis, MO, USA) and nickel-ammonium sulphate (Sigma Aldrich) for up to 20min under visual evaluation. Samples were subsequently washed in Tris-HCl (pH 8.0), PBS and distilled water. Finally, after undergoing ascending dehydration and processing with toluol, samples were coverslipped with Entellan® (Merck, Darmstadt, Germany).

Cell Density and Optical Density Analyses. Individual LGN layers were traced using the polygon drawing tool of Zeiss ZEN 2.0 lite software (Carl Zeiss, Jena, Germany). Immunohistochemical staining of neuronal cell body marker anti-HuC/D showed best differentiation between layers; and was thus used for initial layer tracing and as a template for tracing of the other markers (i.e., anti-MBP and Perls' Prussian blue). Interlaminar koniocellular layers provided visual guidance in separating individual M and P layers. In addition, a previous comprehensive study on LGN laminar arrangements (4) served as anatomical guideline for layer tracing and labeling. Specifically, in posterior parts of the nucleus, the laminar arrangement of the LGN may deviate from the typically described six-layered structure towards a four-layer LGN segment, as also present in the current sample. This four-layer LGN segment comprises two ventral M layers and two dorsal P layers, in which ipsilateral layers P3 & 5 and contralateral layers P4 & 6 are fused in pairs (4).

A stereological analysis of cell density was performed on the immunohistochemical staining of neuronal cell body marker anti-HuC/D. Cell density in each LGN layer was approximated by a cell body count within six uniformly distributed equally sized squares of 60191 $\mu m^2$ in each of the traced LGN layers. This was done using the 'Image Analysis' module, as implemented in the Zeiss ZEN software (version 2.6; Carl Zeiss, Jena, Germany). This procedure resulted in six cell count measures per LGN layer, which were subsequently normalized by area to approximate the cell density in each respective LGN layer.

For the optical density analyses of the anti-MBP and Perls' Prussian blue markers, values of mean intensities were extracted per traced LGN layer through the Zeiss ZEN 2.0 lite software. Normalized optical density measures were then computed by subtraction of the mean intensity from the individual background reference, as measured at an unstained tissue part within a standardized square of 20 $\mu m^2$. This procedure resulted in exactly one normalized optical density measure for each individual LGN layer on the sections stained with anti-MBP or Perls' Prussian blue.

DRAFT

### *In-vivo* MRI

LGN Segmentations on *In-vivo* Quantitative $T_1$ maps. Manual segmentation of bilateral LGNs were performed by two independent raters on the N=27 high-resolution *in-vivo* $qT_1$ maps. In order to standardize the segmentation procedure between raters and across participants, we first computed a histogram of $T_1$ relaxation values (number of bins=1000, bin width=4ms) from each participants' $qT_1$ map. Each histogram was then convolved with a Gaussian filter with sigma=20ms to reduce local signal-to-noise fluctuations. The resulting histograms yielded two clear global peaks, corresponding to the $T_1$ relaxation peaks in gray and white matter in each participant. The $T_1$ relaxation peaks were extracted and subsequently used as windowing parameters for the manual LGN segmentations. Specifically, for each segmentation, the minimum intensity of the $qT_1$ maps was set to the participant-specific white matter $T_1$ relaxation peak, while the maximum intensity was set to the participant-specific gray matter $T_1$ relaxation peak. This was done to optimize the visibility of the LGN and to ensure that all LGN segmentations were based on a similar gray-white matter contrast. The order in which the left or right LGN was segmented was randomized per participant.

Group Template Generation and Population LGN Atlas. To normalize the bilateral LGN masks to a common reference space, we first created a study-specific group template from the N=27 individual high-resolution $qT_1$ maps. The $qT_1$ group template was generated using the *buildtemplateparallel.sh* script implemented in the Advanced Normalization Tools (ANTs, version 2.1.0) software package. The study-specific group template was built in two steps: First, all $qT_1$ images were affine registered using default parameters and averaged to create a globally aligned initial template. This initial template subsequently served as registration target in the first of a total of four iterations of full deformable registration used to create the $qT_1$ group template. Full deformable registration was run with symmetric normalization diffeomorphic image registration (SyN) as transformation model, cross-correlation as similarity metric, and default Gauss regularization [3, 0.5] of the deformation field. Mapping parameters for the SyN transformation model were chosen according to the suggestions in (5). Specifically, the gradient step length was set to 0.5, the number of time discretization points was set to 2, and the integration time step was set to 0.05. Given the high resolution of the qMRI data, the deformable registration was run at five different levels of image resolution (from coarse to fine), with downsampling factors of $2^n$ with n = [4, 3, … 0]. At each image resolution level, the maximum number of iterations was set to $n_{max\_iter}$ = [400, 200, 100, 50, 20]. A total of four iterations of full deformable registration were employed, where each iteration built on the intermediate group template (set as new registration target) and deformed individual data of the previous iteration. The obtained linear and nonlinear registration parameters of the last iteration were then applied to the N=27 $qT_1$ maps using linear interpolation, and all registered images were averaged to create the $qT_1$ group template (Fig. S1A-C and Fig. 3A). Finally, the same registration parameters were applied to the conjoined LGN masks using linear interpolation, followed by averaging of the registered masks within each hemisphere to create a bilateral LGN population atlas (Fig. 3B and Fig. 4A & B).

Cross-Validation of LGN Population Atlas. To assess the prediction accuracy of the bilateral LGN population atlas, we performed a four-fold cross-validation procedure. For cross-validation, four new $qT_1$

DRAFT

group templates were created based on subsets of the data. The four $qT_1$ templates were built in ANTs using the same procedures as described for generating the full N=27 study-specific $qT_1$ group template. Each subset consisted of approximately 75% of the whole $qT_1$ dataset: $N_{foldt1\text{-}3}$=20 participants and $N_{fold4}$=21 participants. For each subset, the corresponding LGN population atlases were additionally computed. The LGN population atlases were subsequently used to predict the anatomical location of the LGN in the remaining participants. Subsets were created pseudo-randomly, such that every participant's LGN was predicted exactly once. Next, the LGN population atlases were warped to the single-subject $qT_1$ maps using SyN in ANTs. For each hemisphere and participant, the registered LGN population atlases were then thresholded in steps of 5% anatomical overlap to most closely match the mean LGN volume across all N=27 participants in the respective hemisphere (for comparison, mean LGN volume left = 113.5mm$^3$, mean LGN volume right = 120.9mm$^3$). Dice coefficients were calculated between the thresholded LGN population atlases and the manually segmented LGN masks in each participant. Finally, Dice coefficients were averaged (weighted by fold size) across participants for each hemisphere. The four-fold cross-validation revealed a good prediction accuracy of the bilateral LGN population atlas on a single-subject level with mean Dice coefficients of 0.80±0.07 and 0.83±0.04 in the left and right hemisphere, respectively.

DRAFT

### SI Figures

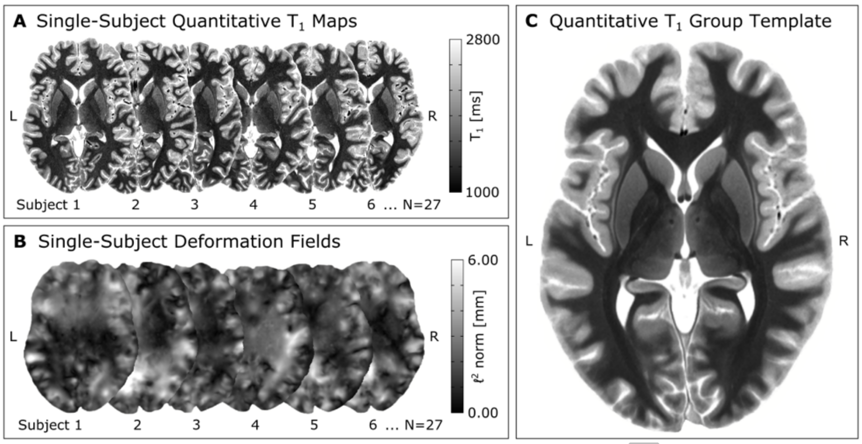


**Fig. S1.** Workflow for creating the study-specific quantitative $qT_1$ group template. **(A)** A total of N=27 single-subject whole-brain $qT_1$ maps served as input for symmetric normalization image registration (SyN) in ANTs. Color scaling of the single-subject $qT_1$ maps indicates the $T_1$ relaxation (in ms) per voxel. **(B)** Four iterations of SyN registration yielded a deformation field for each of the input images, describing the respective voxel displacement for each spatial dimension. For visualization purposes, we here show the Euclidean norm (in mm) of the estimated 3D deformation fields per voxel. **(C)** Following quality control, the deformation fields were applied to the single-subject $qT_1$ maps, and all registered images were averaged to create a study-specific $qT_1$ group template.

DRAFT

**SI References**


1. M. Morawski, *et al.*, Involvement of perineuronal and perisynaptic extracellular matrix in Alzheimer's disease neuropathology. *Brain Pathol.* **22**, 547–561 (2012).
2. C. Stüber, *et al.*, Myelin and iron concentration in the human brain: a quantitative study of MRI contrast. *NeuroImage* **93**, 95–106 (2014).
3. M. Perls, Nachweis von Eisenoxyd in gewissen Pigmenten. *Arch. für Pathol. Anat. und Physiol. und für Klin. Med.* **39**, 42–48 (1867).
4. T. L. Hickey, R. W. Guillery, Variability of laminar patterns in the human lateral geniculate nucleus. *J. Comp. Neurol.* **183**, 221–246 (1979).
5. B. B. Avants, *et al.*, A reproducible evaluation of ANTs similarity metric performance in brain image registration. *NeuroImage* **54**, 2033–2044 (2011).


DRAFT
